# Life history and dispersal timing evolve in response to metapopulation connectedness

**DOI:** 10.1101/476093

**Authors:** Stefano Masier, Dries Bonte

**Affiliations:** Ghent University, Dept. Biology, Terrestrial Ecology Unit, K.L. Ledeganckstraat 35, 9000 Ghent, Belgium. Email: and

## Abstract

Dispersal evolution impacts the fluxes of individuals and hence, connectivity in metapopulations. Connectivity is therefore decoupled from the structural connectedness of the patches within the spatial network. Because of demographic feedbacks, local selection can additionally steer the evolution of other life history traits. We investigated how different levels of connectedness affect dispersal and life history evolution by varying the interpatch distance in replicated experimental metapopulations of the two-spotted spider. We implemented a shuffling treatment to separate local- and metapopulation-level selection.

With lower metapopulation connectedness, an increased starvation resistance and delayed dispersal evolved. Intrinsic growth rates evolved at the local level by transgenerational plasticity or epigenetic processes. Changes in patch connectedness thus induce the genetic and non-genetic evolution of dispersal costs and demographic traits at both the local and metapopulation level. These trait changes are anticipated to impact metapopulations eco-evolutionary dynamics, and hence, the persistence and performance of spatially structured populations.

## Introduction

Habitat fragmentation is, besides habitat loss, one of the most important human-induced drivers of declines in biodiversity (Collinge, 1998; Forman, 2000; Heilman *et al.*, 2002). Most species live in spatially heterogeneous landscapes of suitable habitat patches that are interspersed by unsuitable matrix and connected by dispersal. Connectivity is a species or even population-specific metric that quantifies the fluxes of individuals among patches in the spatial network. It is determined by the individual movement capacities and the number of effective successful dispersers among patches (Tischendorf and Fahrig, 2001). This connectivity is to a large extent determined by the connectedness of the metapopulation. Connectedness defines to which extent a landscape facilitates or impedes the movements of organisms and their genes (Taylor *et al.*, 1993; Rudnick *et al.*, 2012), and refers to its structural properties like the number, shape, dimensions and interdistance– of the suitable patches (Tischendorf and Fahrig, 2000; Wang, Blanchet and Koper, 2014).

Loss of connectivity decreases the chances of reestablishment of extinct populations, thereby putting the metapopulation at risk (Fuller, Doyle and Strayer, 2015; Thompson, Rayfield and Gonzalez, 2017). Connectivity conservation is thus central to metapopulation persistence. While the ecological effects of connectedness loss are well studied from a theoretical (Melián and Bascompte, 2002; Kondoh, 2003; Liao *et al.*, 2017) and empirical perspective (Dobson *et al.*, 2006; Fenoglio *et al.*, 2010; Valladares, Cagnolo and Salvo, 2012), evolutionary consequences are equally anticipated but rarely directly tested (Cheptou *et al.*, 2017). The evolution of inbreeding, mating systems or dispersal rates have been rarely studied so far (Stow *et al.*, 2001; Fahrig, 2003; Andersen, Fog and Damgaard, 2004; Aguilar *et al.*, 2006; Keyghobadi, 2007; Bonte, Masier and Mortier, 2018).

Dispersal is anticipated to be a prime target of selection in metapopulations (Bonte and Dahirel, 2017). It integrates multiple processes during the phases of departure, transience and settlement and should therefore not be treated as a simple trait related to emigration alone (Bowler and Benton, 2005; Clobert *et al.*, 2009). Costs and trade-offs may manifest themselves during each of these three phases (Bonte *et al.*, 2012). Interactions with other life history traits like reproduction or stress resistance (Bonte *et al.*, 2012) give rise to the evolution of life-history syndromes, *i.e.,* consistent correlations of dispersal-related traits (Clobert *et al.*, 2009). These syndromes can manifest themselves at both the individual, population and metapopulation-level (Clobert *et al.*, 2012).

It can be assumed that connectedness loss mostly influences the costs of transfer as movement becomes more costly with increasing distance more distance. These costs are known to lead to the evolution of decreased departure rates (Bonte *et al.*, 2003). Evolutionary adaptations to reduce costs of the other dispersal phases arise as well (Clobert *et al.*, 2012). Habitat connectedness has for instance been shown to be associated with changes in (pre-) departure behaviours (Derr, Aldent and Dinglet, 1981; Bell *et al.*, 2005; Cañete *et al.*, 2007), dispersal timing (Lakovic, Poethke and Hovestadt, 2015), prospecting (Korb and Linsenmair, 2002; Baguette and Van Dyck, 2007) and post-settlement strategies (Vessby and Wiktelius, 2003; Hansson, Bensch and Hasselquist, 2004).

Spatial networks are usually heterogeneous and modular with respect to their topology (Fahrig, 2003; Urban and Skelly, 2006; Ferrari, Lookingbill and Neel, 2007). A few patches are typically more connected to others and thereby serve as hubs in the network (Van Langevelde, 2000; Calabrese and Fagan, 2004). Such asymmetries in connectedness impact local immigration/emigration balances and can lead to source-sink dynamics (Poethke, Dytham and Hovestadt, 2011; Dey, Goswami and Joshi, 2014; Wang, Haegeman and Loreau, 2015). When emigration rates are equal among patches, net immigration rate increase in the most connected patches, thereby elevating local resource competition (Baguette and Schtickzelle, 2003; Amarasekare, 2008; see also box 1). Such local competition can then impose selection on traits related to stress-resistance, or select for a reduced population growth rate to counter resources overshooting in kin-structured populations (McClain, 1995). Finally, if emigration-immigration balances change across patches, different levels of relatedness may affect local reproductive (Macke *et al.*, 2011b) and dispersal strategies (Van Petegem *et al.*, 2018). As a result, even in metapopulations consisting of a network of patches with identical habitat, local selective pressures can be different (Rossetti *et al.*, 2014) and local selection on traits related to density dependence, kin competition and stress resistance may act in concert with metapopulation-level selection on dispersal.

Given the importance of trait evolution for metapopulation dynamics, persistence and rescue, a deep understanding of the eco-evolutionary effects of connectedness changes is essential to correctly predict future population dynamics (Travis *et al.*, 2013; Urban *et al.*, 2016). We engaged in experimental evolution with the spider mite *Tetranychus urticae as* a model (Fronhofer *et al.*, 2014; De Roissart, Wang and Bonte, 2015; Van Petegem *et al.*, 2018), to test whether and how different levels of habitat connectedness affect trait evolution. We particularly focused on traits related to dispersal and reproduction as theory predicts these to be under regional and/or local selection in metapopulations (Duputié and Massol, 2013; Berdahl *et al.*, 2015). We expected individuals in less connected metapopulations to evolve a lower propensity of dispersal, or to evolve lower dispersal costs by means of a higher resistance to the environmental conditions during transfer (e.g., food deprivation). We disrupted putative local selection resulting from systematic changes in densities, and *kin* (genetic relatedness) and *kind* (phenotypic similarity) structure (Van Petegem *et al.*, 2018) by reshuffling mites among local patches in some of the metapopulations. This treatment maintained the distance-related costs of dispersal, and hence, the anticipated metapopulation-level of selection.

## Materials & Methods

### Experimental evolution

Individuals from our focal species, the two-spotted spider mite *Tetranychus urticae* Koch were collected from our stock population (see Supplementary Materials SI1 for a more detailed description) and introduced in 24 artificial habitats composed by a 3-by-3 grid of food patches (bean leaves cuts) connected by Parafilm^©^ bridges (see Fig.1). Due to patch network design, each patch had a different level of connectedness inside the metapopulation (further referred to as *local connectedness*), with central patches having the highest number of links (8), corner patches the lowest (3) and side patches as intermediate (5) value. The bridges varied in length between metapopulations between 4, 8 or 16 cm, thus determining the *metapopulation connectedness* of each setup. Dispersal was considered costly in our experiment both because individuals are not able to forage on Parafilm© and because of the risk of falling from the bridge into the wet cotton and drowning. These transfer costs increase with bridge length.

**Fig. 1:**
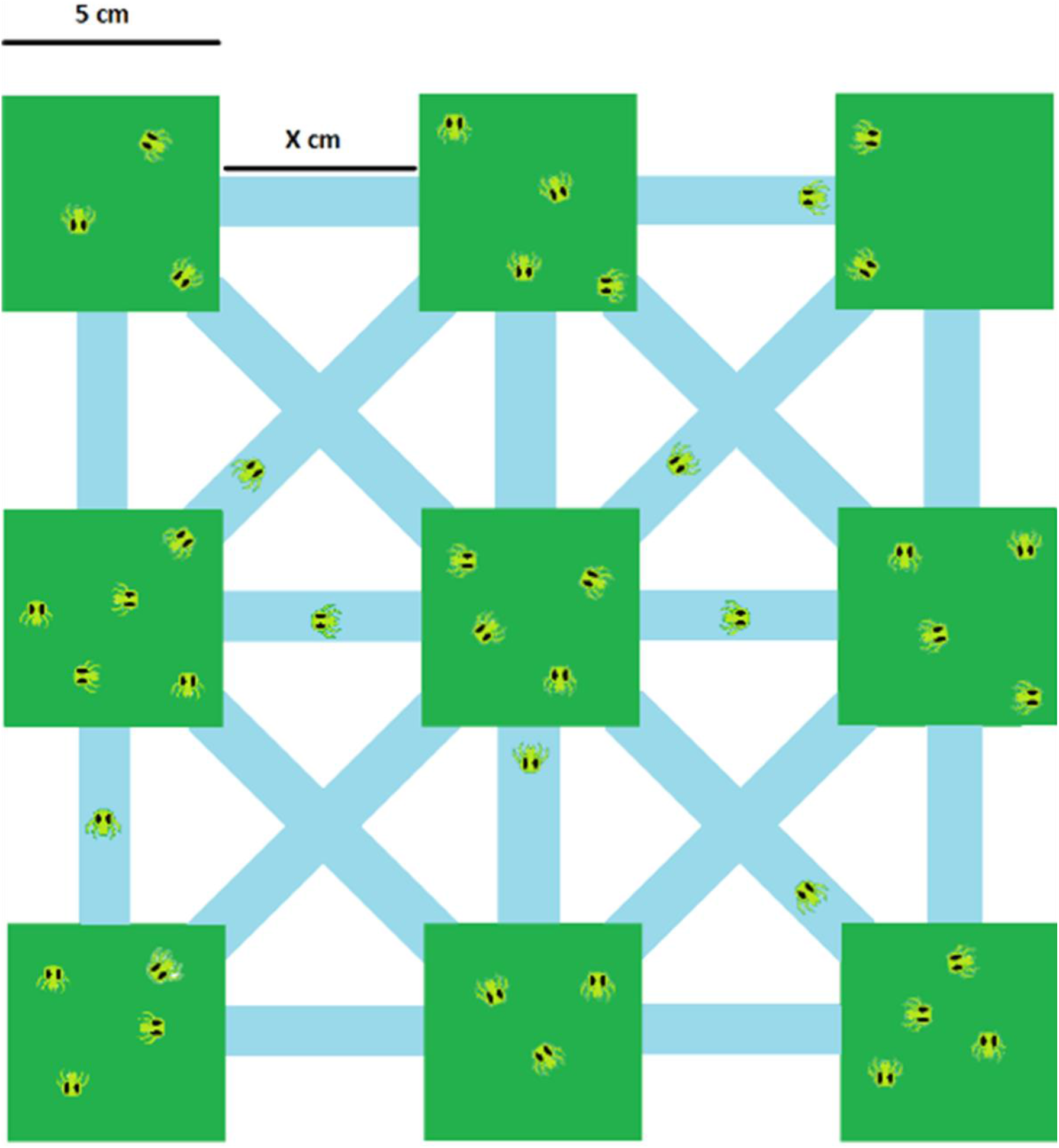
schematic representation of the long-term (6 months) experimental setups. Nine identical patches were arranged in a 3-by-3 grid and connected using Parafilm bridges. The length of the bridge (X cm in the picture) was dependent on the treatment; the diagonal bridges varied depending on the value of X.

Each connectedness level was replicated 8-fold, three of which were subjected to a randomisation procedure of the adult females. We implemented this randomisation procedure to test whether performance would evolve in response to local connectedness as well. In the randomization treatments, all females from a single metapopulation were collected once per week using a thin pen brush, pooled together on a fresh bean leaf to reduce stress as much as possible and then randomly redistributed among the patches of the same metapopulation while maintaining the same local densities as before the collection. This treatment thus destroys any genetic structure of the metapopulation, but maintains the metapopulation demography. Since individuals are dispersing among patches in between the shuffling treatments, they experience identical transfer costs as the non-randomised experimental metapopulations. In order to ensure comparable levels of handling stress between treatments, the females from the non-randomization systems were also collected, not randomized and placed back on their original patch briefly afterwards. We focused on adult females as they are the main dispersal stage for our focal species (Bitume *et al.*, 2011). Contrasting trait values between the randomised and non-randomised experimental metapopulations indicate whether performance evolved at the metapopulation-level (no randomisation effects expected), and/or the local level (randomisation would then remove any signature of local selection).

All experimental metapopulations (24 in total) were initialized in September 2016 with five freshly inseminated females and 2 additional adult males on each patch (hence 1512 individuals overall) using a soft-haired brush and monitored for two weeks to ensure the settlement of the population. This standardised initialisation ensures equal starting conditions and sufficient genetic variation (as for instance demonstrated in Van Petegem *et al.*, 2018). Given the high intrinsic growth rate of the species, populations reached carrying capacity after approximately two generations. After this initialisation period, local population sizes were quantified simultaneously with the reshuffling action, by counting the number of adult females (see De Roissart, Wang and Bonte, 2015). As local changes in population sizes but also their fluctuations are anticipated to drive putative local adaptation in starvation resistance and intrinsic growth, we analysed mean population sizes with GLM having metapopulation and local connectedness as categorical factor and time as a continuous covariate. Experimental metapopulation replicate was modelled as a random factor. Local fluctuations were quantified as their alpha variability, as determined by the coefficient of variance of the local population sizes in time (Wang and Loreau, 2014). These were equally analysed using GLM with experimental metapopulation as a random factor. A more detailed outline of the focal species and the long term experimental setup can be found in Supplementary Materials SI1.

### Trait evolution following common garden

25 weeks from the start of the experiment (approx.12 generations under the experimental conditions; (Hance and Van Impe, 1999), we collected individuals from all the different patches to quantify whether relevant ecological traits had evolved in response to metapopulation connectedness. We quantified performance and dispersal in adult females that were raised for one generation under common garden conditions (see further). We used five adults from each patch in each experimental metapopulation, hence 45 F0 individuals per experimental metapopulation (45 × 24 individuals in total), as the starting material for a generation of F1 females reared under identical conditions. We placed the five females coming from a same patch together into the same common garden and we kept them separated from all the others. F0 females were allowed to lay eggs for two days on a standardised bean leaf, and offspring were subsequently raised under comparable densities (~ 2 individuals / cm^−2^), at 30°C, a 16-8 L/D photoperiod and food *ad libitum*, insured by the low density in relation with the leaf area (~25 cm^2^). This procedure enabled us to study traits independent of plastic responses to the experienced conditions in the metapopulation. After one generation spent in common environment, eventual maternal effects would be mitigated and any difference between lines can be considered due to evolutionary divergence (Macke *et al.*, 2011a).

Given the extent of the trait-quantification experiments, F1 offspring were distributed across experiments to quantify (1) dispersal, (2) starvation resistance and (3) reproductive performance. For reasons of feasibility, we chose to maximise trait quantification across rather than within experimental metapopulations to ensure maximal independent replication: in some cases, we then chose to pool together offspring originating from different patches of the same metapopulation to insure a sufficiently high number of individuals was used.

Below, the structure and statistical analysis for each life-history trait is presented. We present the full models, so including interactions, even when non-significant. We conclude the trait and population analyses with a cross-experimental metapopulation correlation analyses (Pearson’s correlation).

### Dispersal Propensity

This test was performed at metapopulation level. Freshly moulted (1-4 day old) and inseminated females are the dispersing stages in this species (Li and Margolies, 1993, 1994). Such individuals were used for the experiments after rearing one generation under common garden conditions from the same experimental metapopulation replicate. Therefore, data could only be analyzed in function of the metapopulation typology, as this is the anticipated unit of selection. The propensity and the timing of dispersal were tested using a standardized setup of two-patch systems, each composed of two 2.5 × 1.5 cm² patches connected by an 8 cm long, 1 cm wide Parafilm© bridge. On the starting patch, we placed females from F1 at different densities, generating a density gradient between 1.3 and 10.4 individuals/cm². We spanned this range by ensuring equal sample sizes as much as possible around low (5 individuals), intermediate (10) and high (40) densities, the lowest density being similar to what found in nature under a mild infestation scenario (Helle and Sabelis, 1985). A complete set of trials was performed for each metapopulation, with one replica for each density. The two patches were then connected and monitored for four days, as the chance of adult individuals older than five days to disperse is low (Li and Margolies, 1993, 1994). Each day, the number of individuals on the starting and the other leaf was recorded, along with the number of individuals on the bridge at the time of the census. The target leaf to which dispersal took place was refreshed after each count to prevent successfully dispersed females from moving back to the starting patch.

We ran in total 78 two-patches dispersal trials, for a grand total of 1330 monitored individuals. We analyzed both the fraction of emigrants (dispersal rate; individuals leaving the source patch) and the fraction of immigrants (immigration rate; individuals successfully settling on the target patch). These fractions were analyzed using generalized linear mixed model with a binomial error structure and logit-link; metapopulation connectedness, presence/absence of randomization (categorical) and density on the starting patch (continuous), as well as their two- and three-ways interactions were considered as explanatory variables. The metapopulation replicate was added as random effect to control for dependency in responses from individuals originating from the same experimental metapopulation.

Finally, we used the day at which each individual dispersed (dispersal timing) as a dependent variable in a linear model, using again metapopulation connectedness, presence/absence of randomization (categorical) and density on the starting patch (continuous covariate) as explanatory variables, along with their two- and three-ways interactions. We assumed a Poisson distribution for count data with low mean. The metapopulation replicate and dispersal trial replicate were here added as random factors to control for interdependence of data because of shared evolutionary history and experimental setup. Only individuals that performed a dispersal event were included in this analysis so to render this analysis complementary to the emigration rate analysis outlined higher.

### Starvation Resistance

This test was performed at metapopulation level. One experimental arena was prepared for each metapopulation (25 arenas in total: 24 metapopulations + 1 from the stock as external control). Each arena consisted of a 25 cm^2^ square cut from a black plastic sheet mounted on a wet cotton bed into a plastic Petri plate and fixed in position by wet paper strips: the strips partially overlapped with the plastic (~3mm) to form a wet barrier and deter mites from falling into the cotton. For each experimental metapopulation replicate, 5 1-2 day old F1 adult females from the common gardens were placed into an arena. Individuals coming from different patches of the same metapopulation were pooled together as to represent the composition of the original metapopulation. The five out of nine most abundant local populations at the moment of the F0 individuals collection were selected to collect the F1 females. The Petri plates were then stored at room temperature (~25°C).

Based on exploratory trials, we compared the survival percentage after 48h, as this time appeared to be most discriminatory regarding the death/survival balance. Data were analyzed by generalized linear mixed modelling using a binomial distribution, with the connectedness-level and presence/absence of randomization as explanatory categorical variables. As we were only able to have one experimental trial per experimental metapopulation replicate, no random effects were added.

### Reproductive performances

This test was performed at local population and metapopulation level. Single fertilized females from the F1 generation reared under standardised conditions were collected and placed individually on 2.5 × 1.5 cm² bean leaf cuttings, mounted on a bed of wet cotton and kept in place with paper strips. From each experimental metapopulation, 1 or 2 females were collected from each patch, depending on the availability of individuals, for a grand total of 370 females across the 24 metapopulation replicates. The leaves were placed in a temperature regulated cabinet at 30°C for ten days with a 16-8 L/D photoperiod. They were daily moisturized to prevent desiccation of the leaves and the escape of the mites through the cotton layer. Each female was monitored daily, recording her status (dead or alive), as well as the number of eggs, larvae and juveniles present on the patch. The number of individuals from these three age classes were subsequently summed as a measure of reproductive performance. We ran a GLM assuming a quasi-Poisson distribution for the count data, to correct for overdispersion (Cameron and Trivedi, 1990). Reproductive performance at day 6 was used as the dependent variable to avoid effects of overlapping generations, as the following generation F2 individuals did not yet reach adulthood at the time. As reproductive performance was hypothesised to evolve in relation to local conditions as well, we added patch location of the F0 females (centre, corner or edge) as an independent categorical variable in addition to the earlier used metapopulation-level connectedness and randomisation treatments. The experimental metapopulation replicate was added as random effect to control for shared evolutionary history. As the reproductive rate was calculated at patch or treatment level but not at the individual level, females that died before the end of the test were retained in the analysis as their performances still impacted the reproductive success of their (meta-)population of origin. A complementary analysis was performed to check if the day of death was significantly different between treatments, using the same model structure as previously described.

## RESULTS

### A. Local population dynamics

Results from the statistical analyses are given in Supplementary Materials SI2. From the period after initiation of the setup, and hence, after the first month of population growth, no differences in local population sizes or local fluctuation here-in were recorded. Population sizes were equally stable in time, indicating the experimental metapopulations were at equilibrium. Contrary to expectations, no differences according to local connectedness were detected. Analyses over the last four weeks (~2 generations) before the trait assessment revealed neither significant differences.

### B. Trait evolution

#### 1. Dispersal

Evolved emigration and immigration rate (success rate) were not dependent on the connectedness or randomization treatment; only the starting density had a significant and positive (see Fig SI3.1 in Supplementary Materials SI3) impact on the emigration rates (Table 1). Individuals that evolved in the metapopulations with the lowest level of connectedness showed overall delayed timing of dispersal (day of departure) (Figure 2a), but this effect was especially pronounced when no randomization was applied (significant interaction randomization treatment x metapopulation level of connectedness; Table 1).

**Fig. 2a (top panel):**
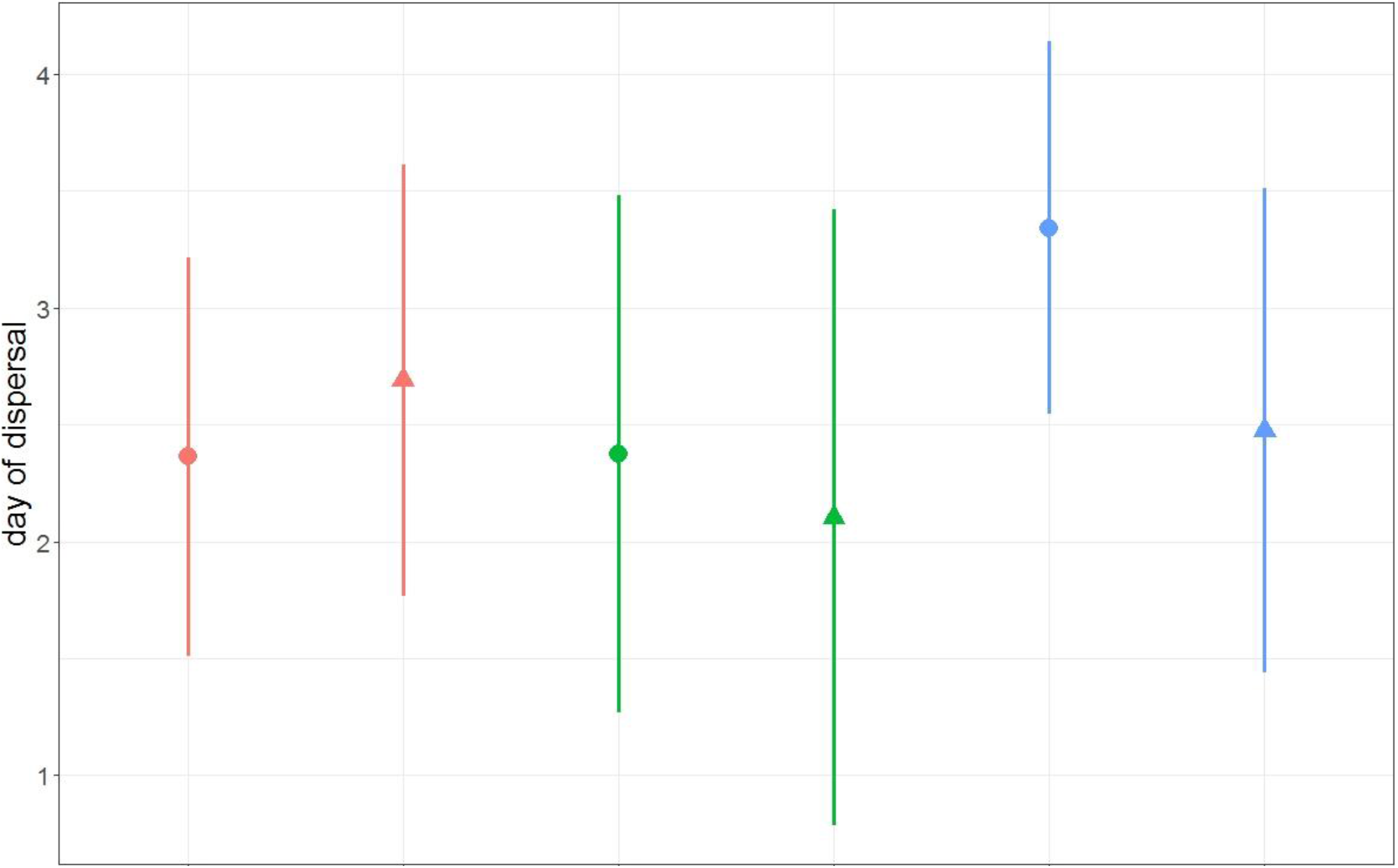
the effect of connectedness on the individual day of dispersal in the different treatments (means ± sd).

**Table 1:** Full Model Analysis of Deviance table (Type III Wald *χ*^2^-tests) for emigration and immigration rate and dispersal timing in relation to the connectedness, randomization of the metapopulation in which individuals evolved and density in the dispersal trial.

| Dependent variable | Explanatory variable | df | $\chi^2$ | Pr(>$\chi^2$) |
| --- | --- | --- | --- | --- |
| Emigration rate | Metapopulation Connectedness | 2 | 1.5251 | 0.6765 |
|  | Randomization | 1 | 0.0172 | 0.8957 |
|  | Starting Density | 1 | 7.96 | 0.0048 (**) |
|  | Metapopulation Connectedness * Randomization | 2 | 0.1978 | 0.9058 |
|  | Metapopulation Connectedness * Starting Density | 2 | 2.0713 | 0.3549 |
|  | Randomization * Starting Density | 1 | 0.3847 | 0.5351 |
|  | Metapopulation Connectedness * Randomization * Starting Density | 2 | 3.5738 | 0.1675 |
| Immigration rate | Metapopulation Connectedness | 2 | 0.6346 | 0.8885 |
|  | Randomization | 1 | 0.5902 | 0.4423 |
|  | Starting Density | 1 | 0.3421 | 0.5586 |
|  | Metapopulation Connectedness * Randomization | 2 | 0.6865 | 0.7095 |
|  | Metapopulation Connectedness * Starting Density | 2 | 1.3771 | 0.5023 |
|  | Metapopulation Randomization * Starting Density | 1 | 0.0099 | 0.9208 |
|  | Metapopulation Connectedness * Randomization * Starting Density | 2 | 1.1632 | 0.5590 |
| Individual dispersal day | Metapopulation Connectedness | 2 | 17.1631 | 0.0001875 |
|  | Randomization | 1 | 1.7121 | 0.1907190 |
|  | Starting Density | 1 | 0.0001 | 0.9916746 |
|  | Metapopulation Connectedness * Randomization | 2 | 7.0688 | 0.0291764 (*) |
|  | Metapopulation Connectedness * Starting Density | 2 | 0.7331 | 0.6931065 |
|  | Randomization * Starting Density | 1 | 0.5802 | 0.4462486 |
|  | Metapopulation Connectedness * Randomization * Starting Density | 2 | 4.4365 | 0.1087995 |

#### 2. Starvation resistance

The proportion of individuals surviving 48 hours of starvation only depended on the connectedness experienced during experimental evolution, with highest survival rates for individuals that evolved in the least connected metapopulation relative to the two other treatments (Analysis of Deviance, Type III test: *χ*^2^=11.89, P=0.003; Fig. 2b). Survival rates did not differ between the randomization procedures (Analysis of Deviance, Type III test: *χ*^2^=0.12, P=0.720) nor on the interaction between both (Analysis of Deviance, Type III test: *χ*^2^=0.85, P=0.651).

**Fig.2b (bottom panel):**
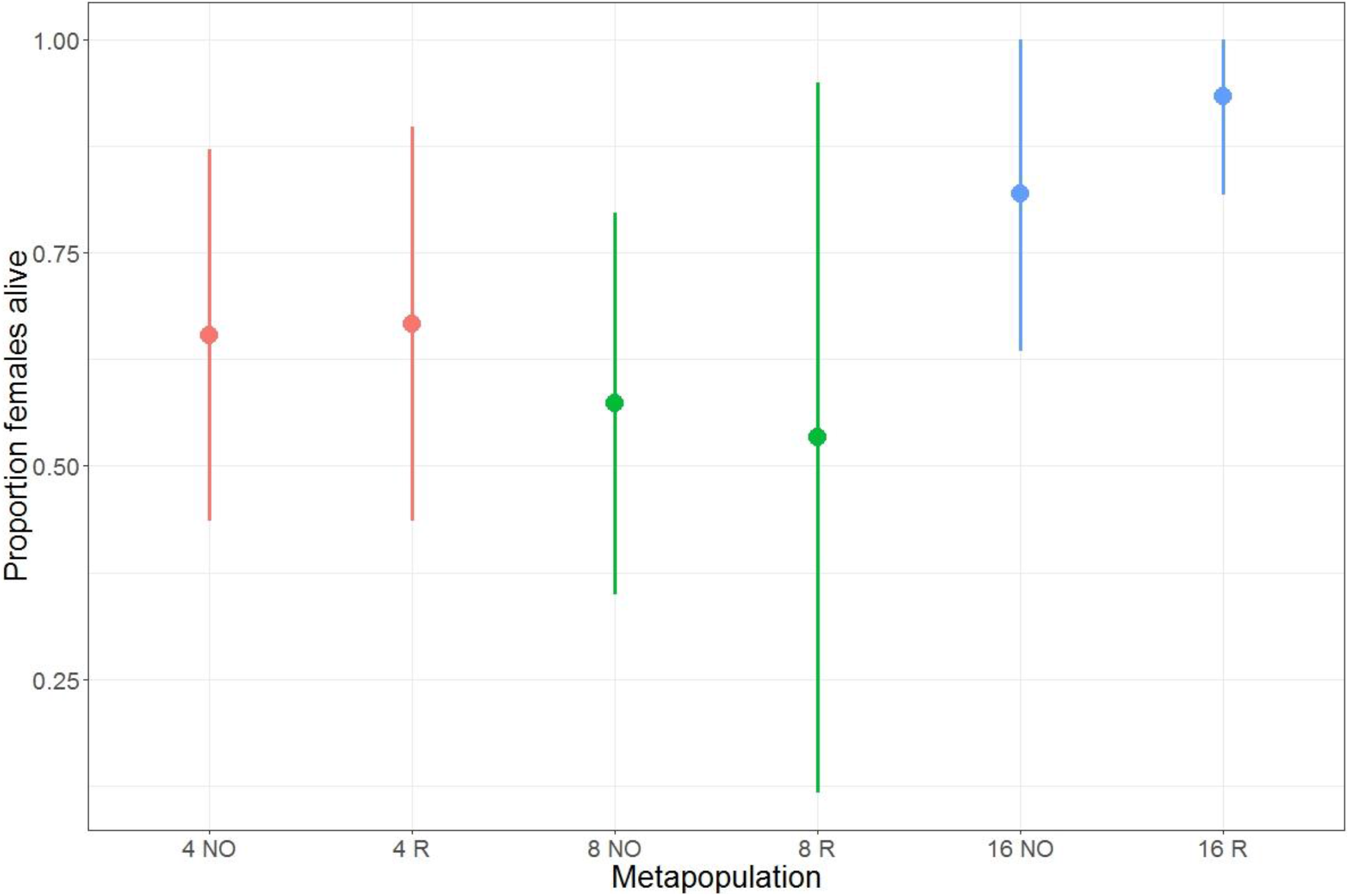
proportion of alive individuals (means ± sd) after 48h depending on the treatment of the source metapopulation.

#### 3. Reproductive performance

The number of the females that died during the test did not differ among treatments (see Supplementary Materials SI4). The local connectedness of the patch the individuals originated from affected reproduction significantly (Analysis of Deviance, Type III test: *χ*^2^=9.6736, P=0.008, Fig. 3). In general, reproductive performance was overall the lowest in offspring from individuals that were collected in the central patch (Posthoc Tukey test: corner-center contrast: t-ratio = −2.895, P=0.0112; edge-center contrast: t-ratio = −3.073; P=0.0065). Reproductive performance was not affected by the metapopulation connectedness or the randomization procedure (see table SI4.2 in Supplementary Materials SI4 for the complete analysis).

**Fig. 3:**
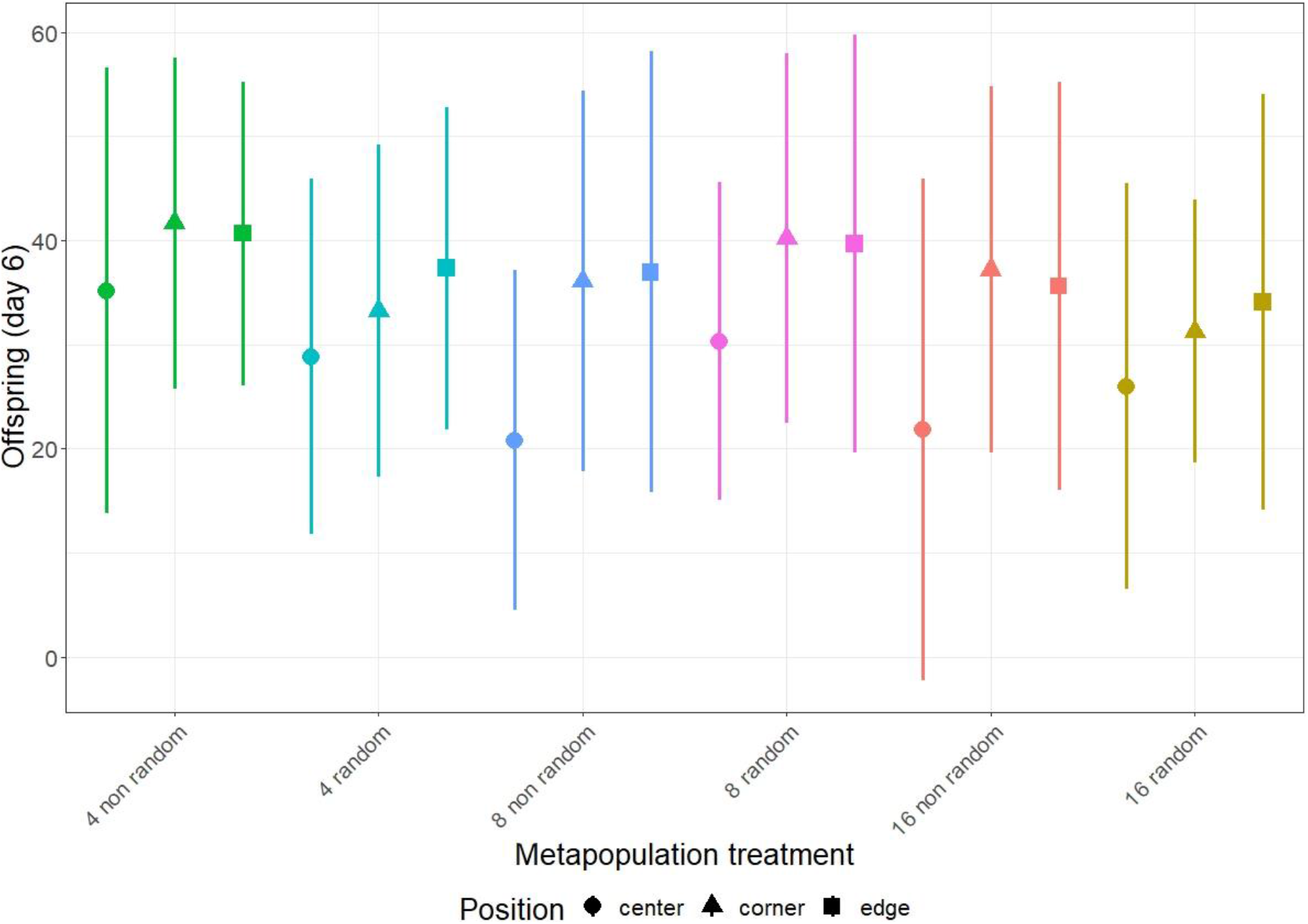
Reproductive performance measured as the number of offspring after 6 days (mean ± sd), for each combination of fragmentation treatment and relative position of the patch of origin of the individuals (before common garden period).

### C. Trait correlations at the metapopulation-level

Across experimental metapopulations, dispersal latency is positively correlated with reproductive performance (*r*23=0.52; p=0.012). Populations that evolved a high intrinsic growth rate thus evolved a delayed dispersal. Metapopulations where indivuals evolved a delayed dispersal attained larger local population sizes during the 6 months of the experimental evolution (*r*23=0.43, p=0.043) (Figure 4 and Supplementary Materials SI5 for raw correlation coefficients and significance levels).

**Fig.4:**
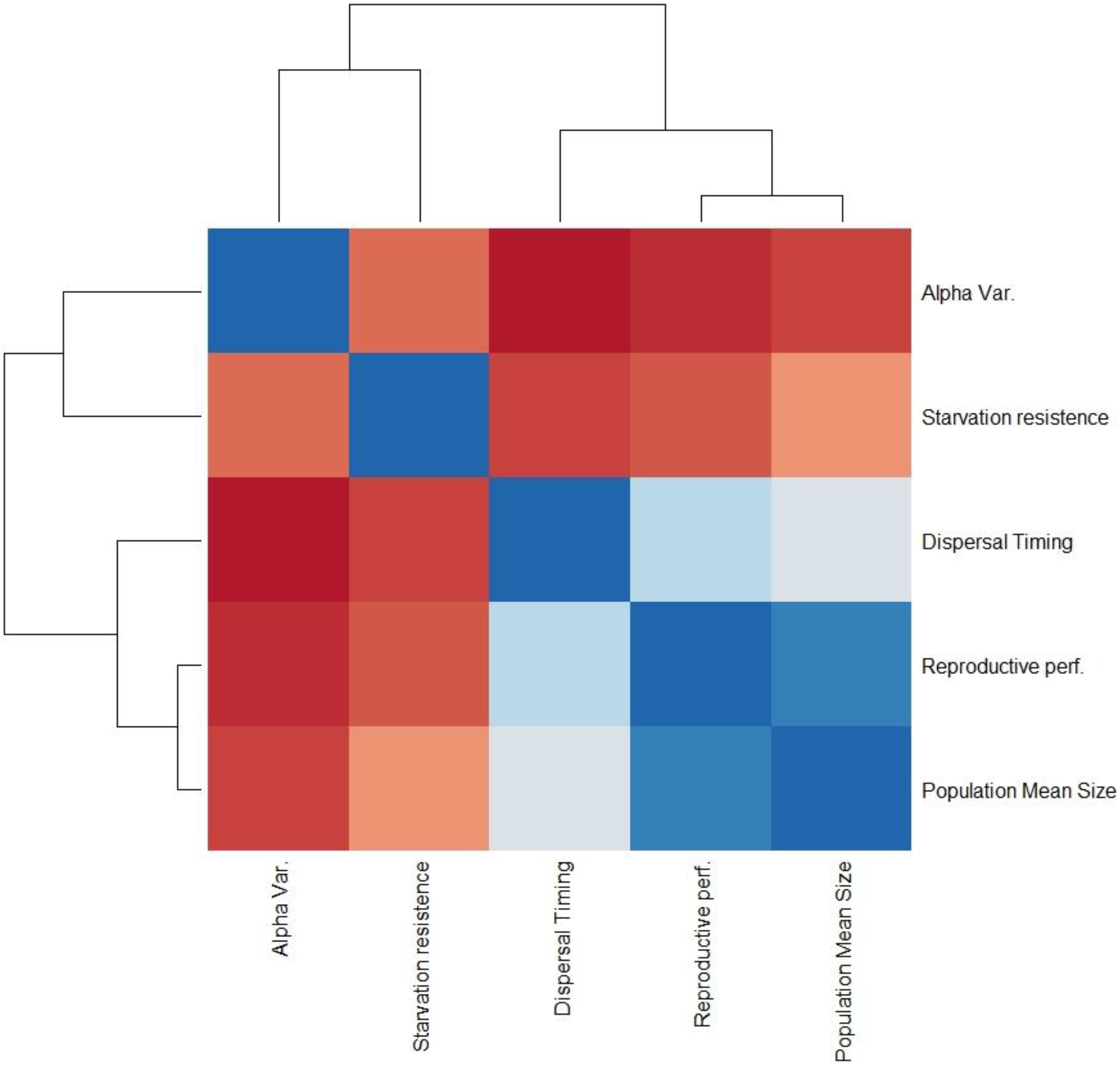
correlations among traits, population size and variance as attained during the experimental evolution setup. Darker colors indicate higher values (in module) of Pearson’s coefficients. Blue squares represent positive relationships; red squares are negative. For the exact values of the coefficients and the complete significance table, please see Supplementary Materials SI5.

## DISCUSSION

Dispersal is a complex life history trait encompassing multiple, often tightly coupled stages. To date, most evidence for dispersal evolution in metapopulations stems from studies on emigration rate (Olivieri, Michalakis and Gouyon, 1995; Mathias, Kisdi and Olivieri, 2001; Gyllenberg, Parvinen and Dieckmann, 2002; Ronce, 2007). However, simultaneous evolution of traits related to the multiple dispersal stages, including transience and settlement, has been theoretically demonstrated to lead to often complex feedbacks which may or may not impact the emigration rate (Travis *et al.*, 2013; Delgado *et al.*, 2014). Our experimental work demonstrates that changes in metapopulation connectedness altered the joint evolution of dispersal timing, dispersal costs and reproduction. Connectedness loss did–contrary to expectations- not lead to the evolution of decreased emigration rates *per se.*

The evolution of dispersal timing has been theoretically addressed in response to either dispersal at adulthood (breeding dispersal) or at birth (natal dispersal) (Johst and Brandl, 1999; Hirota, 2004; Lakovic, Poethke and Hovestadt, 2015; Lakovic, Mitesser and Hovestadt, 2017), and only recent studies have started addressing the issue at a more detailed time scale (Li and Kokko, 2019).

Here, we found evolved breeding dispersal to be delayed in the least connected metapopulations while no effect of density was found. This delayed dispersal is relevant for connectivity as most spider mites disperse when freshly moulted, so up to two days after development into the adult instar (Hussey and Parr, 1963; Li and Margolies, 1993). Interestingly, the delay was especially pronounced in the non-randomized treatment, indicating that evolution in response to local kin or kind interactions, or other local selection pressures is a more prominent driver than metapopulation-level selection from changes in connectedness. Delayed dispersal can be mechanistically interpreted as a delayed density dependence as individuals delaying dispersal are more likely to be longer engaged in local competitive interactions. In this sense, it reflects density-dependent emigration strategies (Saether, Engen and Lande, 1999; Hovestadt and Poethke, 2006) with density thresholds evolving to higher levels in disconnected metapopulations.

Individuals from the least connected metapopulation also evolved the largest starvation resistance, but independent of the randomization treatment. As we found no correlation between evolved starvation resistance and dispersal timing across metapopulations, both traits evolved jointly but independently in the non-randomized metapopulations. This accords with the perspective brought forward by Bonte & Dahirel (2017) that dispersal needs to be treated as a central and independent trait in life history evolution. Given the lack of a consistent correlation between dispersal timing, local mean population sizes and their variability, we do not attribute the evolution of starvation to result from adaptations to local densities and putative changes in local competition. Rather, we explain the evolution of starvation resistance as a response to metapopulation-level costs of dispersal, i.e. the energetic and risk costs during transfer on the bridges (Bonte *et al.*, 2012). Hence, an “endurance-oriented” strategy evolved in the least connected metapopulations. Taken together, the evolution of delayed dispersal and of the costs of dispersal should equalize connectivity as measured by long-term immigration rates among the different metapopulations across the different levels of habitat connectedness (Bélisle, 2005; Baguette and Van Dyck, 2007). We here thus demonstrate the prevalence of a predicted but not yet documented eco-evolutionary feedbacks in metapopulations (Bonte, Masier and Mortier, 2018).

Building up energy reserves to survive long periods of food shortage is a costly process that allocates resources from other vital processes. Starvation resistance is commonly associated with larger body mass, and usually has consequences on fertility, reproduction timing and longevity (Tessier *et al.*, 1983; Hoffmann and Harshman, 1999; Rion and Kawecki, 2007). However, we did not detect such trade-offs as reproductive performance only diverged in response to the local patch type and not in relation to the connectedness of the metapopulation in which they evolved. The systematic lower reproductive performances in individuals from central patches in the network is intriguing as patches were identical in terms of quality and size throughout the network in both the unmanipulated metapopulations (natural dispersal) and the ones that are shuffled. Reduced reproductive performance is expected to evolve in response to intense competition (Krüger, Liversidge and Lindström, 2002; Agrawal, Underwood and Stinchcombe, 2004) and patches with higher local connectedness should theoretically receive more individuals than those with fewer connections, independent of the metapopulation-level connectivity. We could, however, not attribute variation in reproduction to selection from local competition as both mean and variance of local population sizes did not diverged significantly in response to local and metapopulation-level connectedness.

Even more paradoxically, this signature was not destroyed by randomization. This implies that the observed lower reproductive performance, as measured after one generation of common garden, must originate from persisting intergenerational plasticity rather than from genetic evolution. A recent meta-analysis indeed demonstrated that maternal effects are an influential source of phenotypic variation across the tree of life (Moore, Whiteman and Martin, 2019). Such transgenerational plasticity is common in *T. urticae* (Bitume *et al.*, 2011; Magalhães *et al.*, 2014; Marinosci *et al.*, 2015; Van Petegem *et al.*, 2015), and even grandmaternal effects have been recorded in dispersal-related traits (Bitume *et al.*, 2014). As reproductive performance was positively correlated across metapopulations with dispersal latency, local condition thus affect body condition, dispersal timing and reproduction across generations in a tangled way. We were unfortunately not able to study variation in dispersal in relation to local connectedness, so whether local changes in kin or kind structure are driving these transgenerational responses remains unclear and deserver further study, also from a general perspective. Overall, this insight puts experimental common-garden procedures at question to separate plastic from genetic effects in trait expression. As even conditions experienced by grand-mothers have the potential to impact life history traits in complex manners, common garden experiments as performed by us and many others (see De Meester *et al.*, 2019) need to be interpreted with the needed precaution in terms of the exact underlying mechanism. Ideally they are accompanied by genomic analyses if the loci underlying trait expression are known (for dispersal some candidate genes have been detected; Saastamoinen *et al.*, 2018). From an ecological perspective, these insights remain nevertheless important as transgenerational effects may much faster than genetic selection feedback on the ecological dynamics (Galloway, 2005; Drummond and Ancona, 2015) and leave signals that are currently interpreted as feedbacks from genetic evolution (De Meester *et al.*, 2019).

We demonstrate that evolutionary dynamics in metapopulations do not follow simple theoretical predictions based on dispersal and genetic selection alone. Local and metapopulation-level dynamics interact with each other and influence the adaptation of the individuals at both levels, with ecological population dynamics influencing evolutionary dynamics and the other way around (Hanski, 2012). Stress resistance, for instance, which we found to evolve in response to habitat fragmentation, can potentially turn into pre-adaptation to other kind of stressors, i.e. suboptimal host plants or novel habitats and enhance persistence under environmental change (Jenkins, Chaisson and Matin, 1990; De Roissart *et al.*, 2016).

Habitat fragmentation is a central problem in conservation biology as it is expected to reduce connectivity and eventually lead to species loss and disintegration of local food webs (Thompson, Rayfield and Gonzalez, 2017). However, connectivity between patches can be evolutionary rescued through adaptation to higher dispersal costs or development of novel dispersal mechanisms (Kendrick *et al.*, 2017). Given that evolution of dispersal-related traits shifts cost-benefit balances towards a more beneficial ratio in metapopulations (Bonte *et al.*, 2012), these evolutionary dynamics adaptations will feedback on other traits (e.g. different allocation of energy reserves to reproduction) as well and thereby impact the ecological processes (Bonte, Masier and Mortier, 2018). Such an integrated view on dispersal, life history evolution and costs shifts in response to habitat fragmentation is currently lacking (Cheptou *et al.*, 2017) but highly needed to advance the predictive ecology of species and ecosystems, especially in the fast-changing world of today (Urban *et al.*, 2016). While theory focuses, for reasons of tractability, on simple dynamics (Govaert *et al.*, 2019), we are able to demonstrate more realistic multivariate life history divergence in response to changes in habitat connectedness as caused by extended evolutionary processes (Laland *et al.*, 2015; Futuyma, 2017).

The decoupling of habitat connectedness and population connectivity could impact predictions from theory that assumes a direct and positive link between dispersal and habitat connectedness (Legrand *et al.*, 2017). The evolution of often-overlooked traits like dispersal timing, stress resistance or individual growth rates rather than dispersal rates *per se* could then act as a rescue mechanism for fragmented populations through their impact on metapopulation variability, local stabilizing and spatial synchronizing effects (Wang, Haegeman and Loreau, 2015).

### Box 1: Asymmetric immigration rates

Theoretical demonstration how differences in local connectedness impose asymmetric immigration rates, and thus mean carrying capacity. In a spatial network as implemented in our experimental evolution, we assume local resource availability determining local carrying capacity *K*, and dispersal rates *d*. If we assume the population to be regulated at *K*, then *dK* individuals are emigrating from each patch, and those emigrants will be equally divided across the number of connections to the other patches *C*. Given a certain dispersal mortality *μ*, each ‘target’ patch will then receive (1-*μ*)*(d*\*K/C*) immigrants. If dispersal is occurring after local density regulation (as expected in our experiments because of female dispersal), then the eventual population size will be largest on the most connective patches and reaching a new equilibrium

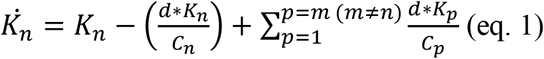

This yields for the different local patches to the following 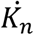:

| PATCH | EMIGRANTS (PER BRIDGE) | IMMIGRANTS |
| --- | --- | --- |
| **Centre** | $(d*K/8)$ | $4(d/3)+4(d/5) = (32/15)d \approx \mathbf{2.13d}$ |
| **Corner** | $(d*K/3)$ | $2(d/5)+ d/8 = (21/40)d \approx \mathbf{0.52d}$ |
| **Side** | $(d*K/5)$ | $2(d/3)+2(d/5)+ d/8 = (143/120)d \approx \mathbf{1.19d}$ |

