## Supplementary materials for "Life history and dispersal timing evolve in response to metapopulation connectedness"

**Supplementary Information 1: experimental setup**

*Species description and stock characteristics*

Our study species is the haplodiploid spider mite *Tetranychus urticae* Koch (Acarina: Tetranychidae), one of the most important herbivore pests worldwide and a commonly used model species (Grbic et al., 2007). The species has a generation time of approximately 15 days at 30 °C (Hance & Impe, 1999), and its fecundity is high (Tehri, 2014).

The stock “LS-VL” population was initiated from wild-caught mites from the Ghent University botanical garden in Belgium in October 2000 (Cazaux et al., 2014; Van Leeuwen et al., 2008), and maintained on full bean plants (*P. vulgaris* L., variety ‘Prelude’) since then. The plants were raised in walk-in climatically controlled rooms and transferred to the stock population, which was kept in a separate room to avoid contamination. The mites were maintained on full plants, inside top-open plastic crates and under a 16-8 L/D regime at a constant temperature of ~25°C. The average size of the stock population is measured in the thousands, with the chance of reaching even higher numbers by providing extra food for ~1 month. Every two/three years, some individuals from outside the stock population (usually collected in the same spot the stock population originated from) were added to maintain genetic variability.

*Experimental evolution*

We developed experimental metapopulations, with each setup composed of nine habitat patches organized in a three-by-three grid. Each patch consisted of a square (5 cm × 5 cm) cut from a bean leaf collected from ~2 weeks old plants. The patches were connected to each other by Parafilm bridges: each bridge was 0.5 cm wide, while the length varied according to the connectedness treatment (see further). The leaves and bridges were mounted on a bed of wet cotton, to keep them hydrated and to prevent mites from escaping or dispersing outside the designed pathways. Each experimental metapopulation was separately kept in a Vaseline-coated, open plastic box to further reduce the risk of mites escaping. Distilled water was added daily to the cotton to keep moisture levels constant. The boxes were placed inside two climatically controlled walk-in rooms, with a constant temperature (25°C) and a 16-8 LD photoperiod for the whole duration of the experiment. To exclude any possible effect of location within the room, the position of all boxes was randomized once per week.

The “metapopulation connectedness” treatment consisted of 3-by-3 grids of patches connected by bridges of 4, 8, or 16 cm of length. Diagonal bridges, whose length was dependant on the grid bridges length, were added to connect the central and corner patches and the four lateral patches (see fig.1, main text).

Although individual mites are able to walk distances up to several meters per day (Krainacker & Carey, 1990), these lengths were chosen because the lengths of 8 or 16 cm were demonstrated to constrain dispersal and impose selection due to reduced feeding time and distance-dependent mortality costs of ending up in the cotton (Bitume et al., 2013 and Masier et al., unpub data). The total area of leaves and, hence, resource availability was identical across all the connectedness treatments. As a result of the patch network design, each patch had a different level of connectedness inside the metapopulation (further referred to as *local connectedness*), with central patches having the highest (connected to every other patch), corner patches the lowest (connected only to three other patches) and side patches as intermediate (connected to five patches) value.

All patches in each experimental metapopulation were replaced weekly on the same day with new freshly-cut bean leaf squares. The adult females, randomized or not, were directly moved onto the new leaves as described higher. Old leaves were then laid upon the new ones, supported by toothpicks to avoid mould spread, and left in position for 48h before being disposed of. This was done to allow larvae, juveniles and adult males left on the old leaves to move to the new ones, as well as to provide some time for the eggs laid on the old leaf to hatch (Praslicka & Huszár, 1989)

**Supplementary Information 2: local population dynamics**

1. ***Long-term population dynamics***

   *Fig.SI2.1: local population size (upper panel) and alpha variability (lower panel) (mean ± sd) over 6 months of experimental evolution, depending on metapopulation treatment and position*

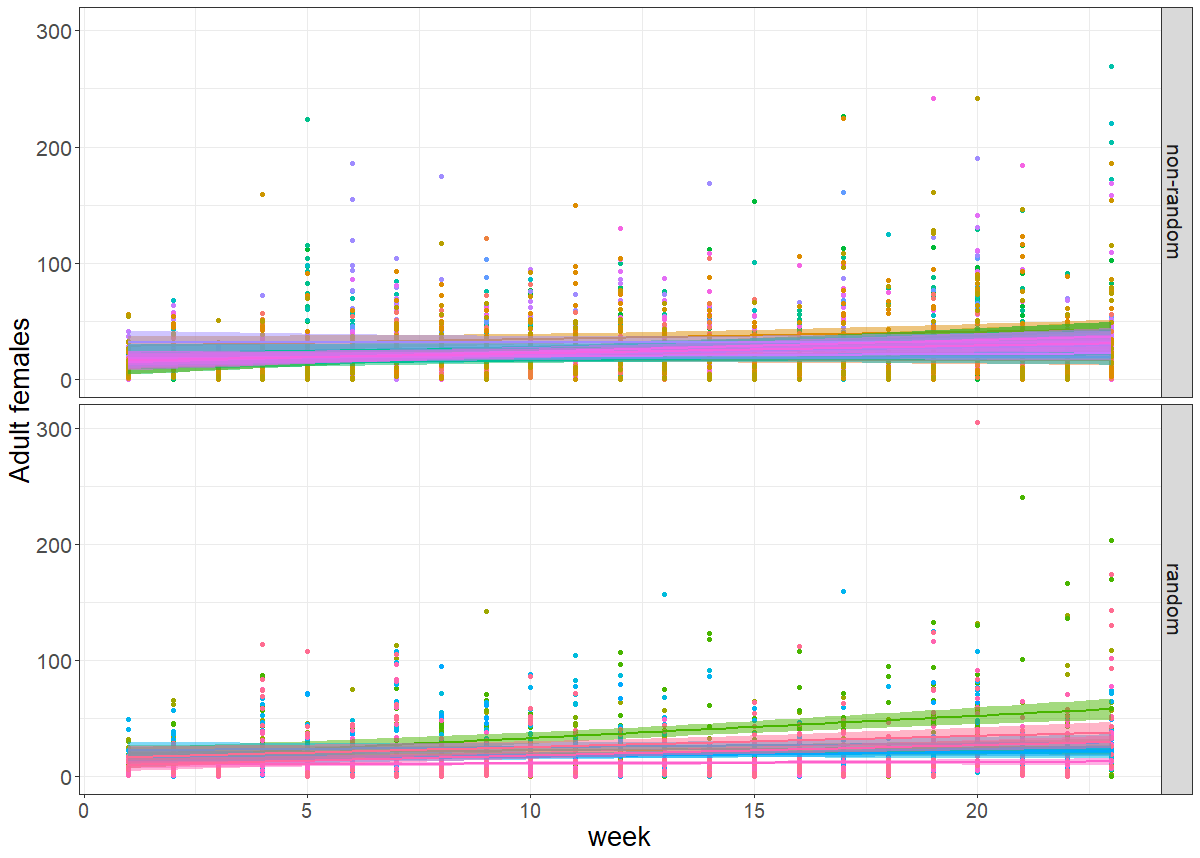

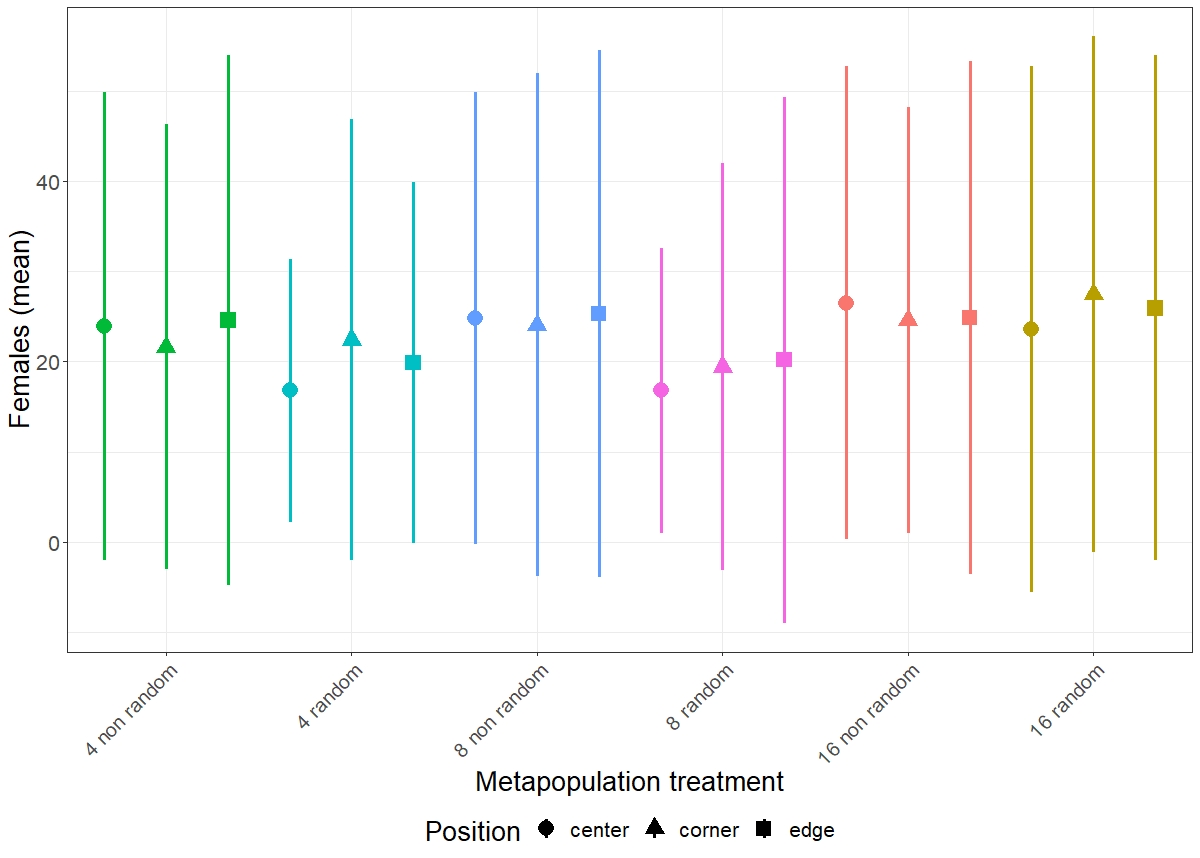

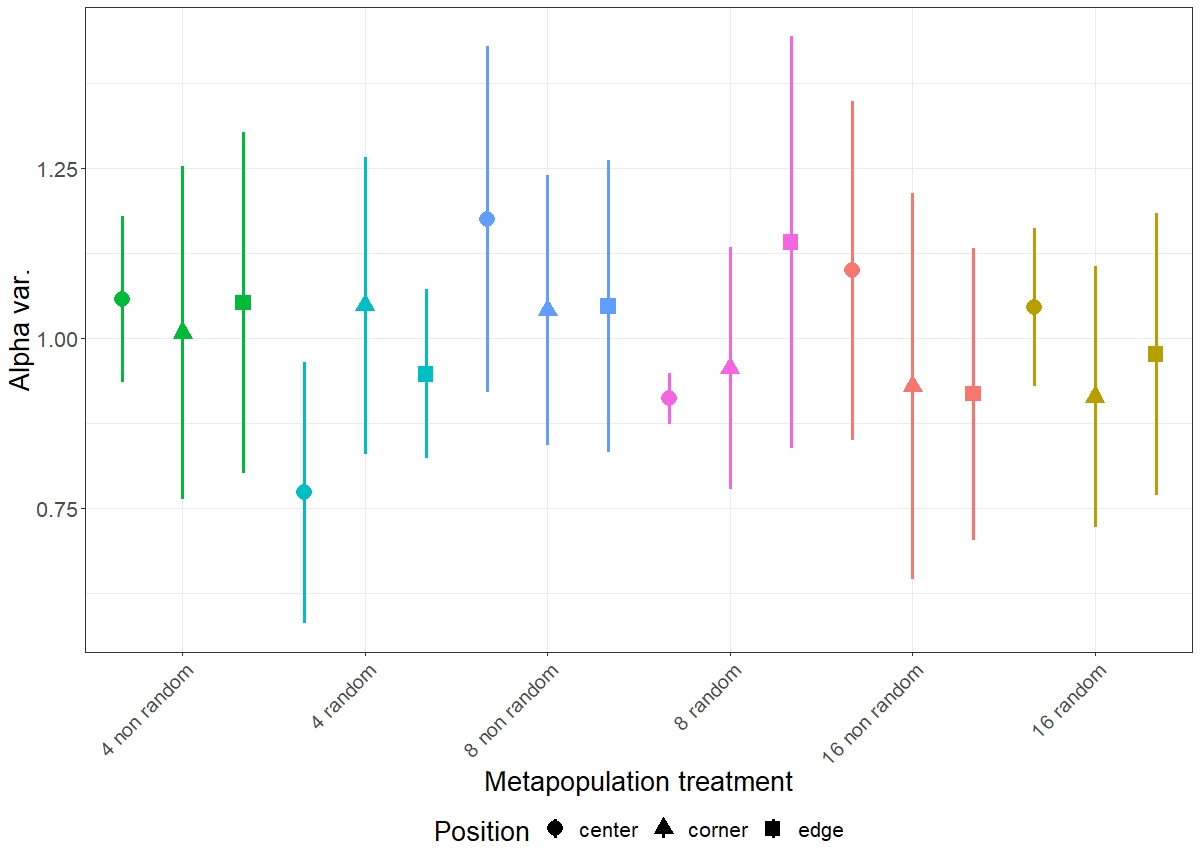

*Fig.SI2.2: metapopulation size, calculated using the number of adult females as proxy, and linear regression line for each metapopulation during the 6 months of the experimental evolution, divided by randomization treatment.*

***Table SI2.1a***

| **Explanatory variable** | **Χ^2^** | **Df** | **Pr(>Χ^2^)** |
| --- | --- | --- | --- |
| Time | 1.7447 | 1 | 0.1865 |
| Global connectedness | 0.0025 | 2 | 0.9987 |
| Randomization | 0.0237 | 1 | 0.8776 |
| Local connectedness | 0.9159 | 2 | 0.6326 |
| Global connectedness * Randomization | 0.8134 | 2 | 0.6658 |
| Time*Global connectedness | 0.6595 | 2 | 0.7191 |
| Time*Randomization | 0.1508 | 1 | 0.6978 |
| Time*Local connectedness | 0.0623 | 2 | 0.9693 |
| Global connectedness*Local connectedness | 0.5920 | 4 | 0.9640 |
| Randomization*Local connectedness | 0.0393 | 2 | 0.9806 |
| Time*Global connectedness*Randomization | 0.1202 | 2 | 0.9416 |
| Global connectedness*Randomization*Local connectedness | 0.9548 | 4 | 0.9166 |
| Time*Global connectedness*Local connectedness | 1.2520 | 4 | 0.8695 |
| Time*Randomization*Local connectedness | 0.0113 | 2 | 0.9944 |
| Time*Global connectedness*Randomization*Local connectedness | 1.6805 | 4 | 0.7943 |

***Table SI2.1b***

| **Explanatory variable** | **Χ^2^** | **Df** | **Pr(>Χ^2^)** |
| --- | --- | --- | --- |
| Global connectedness | 0.5028 | 2 | 0.77770 |
| Randomization | 2.9091 | 1 | 0.08808 (.) |
| Local connectedness | 0.4591 | 2 | 0.79491 |
| Global connectedness * Randomization | 1.1213 | 2 | 0.57083 |
| Global connectedness*Local connectedness | 1.5756 | 4 | 0.81317 |
| Randomization*Local connectedness | 4.2834 | 2 | 0.11746 |
| Global connectedness*Randomization*Local connectedness | 5.6560 | 4 | 0.22636 |

*Table SI2.1: ANOVA (Analysis of Variance: Wald Type-III chisquare test) results for mean (a) and variance (b) of the local population sizes over the 6 months of the experimental evolution.*

2. ***Short-term (4 weeks) population dynamics*** *Fig.SI2.3: local population size (upper panel) and alpha variability (lower panel) (mean ± sd) over the last 4 weeks (~2 generations) of the experimental evolution, depending on metapopulation treatment and position*

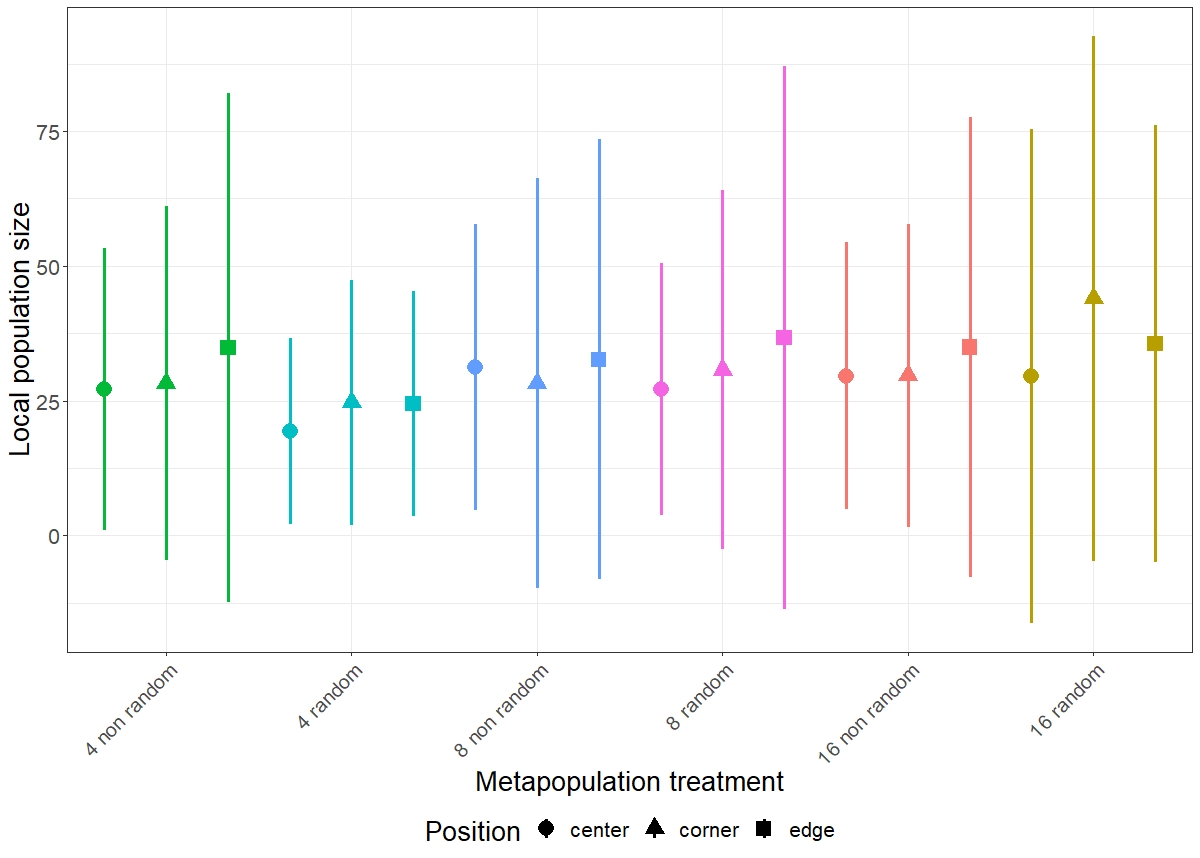

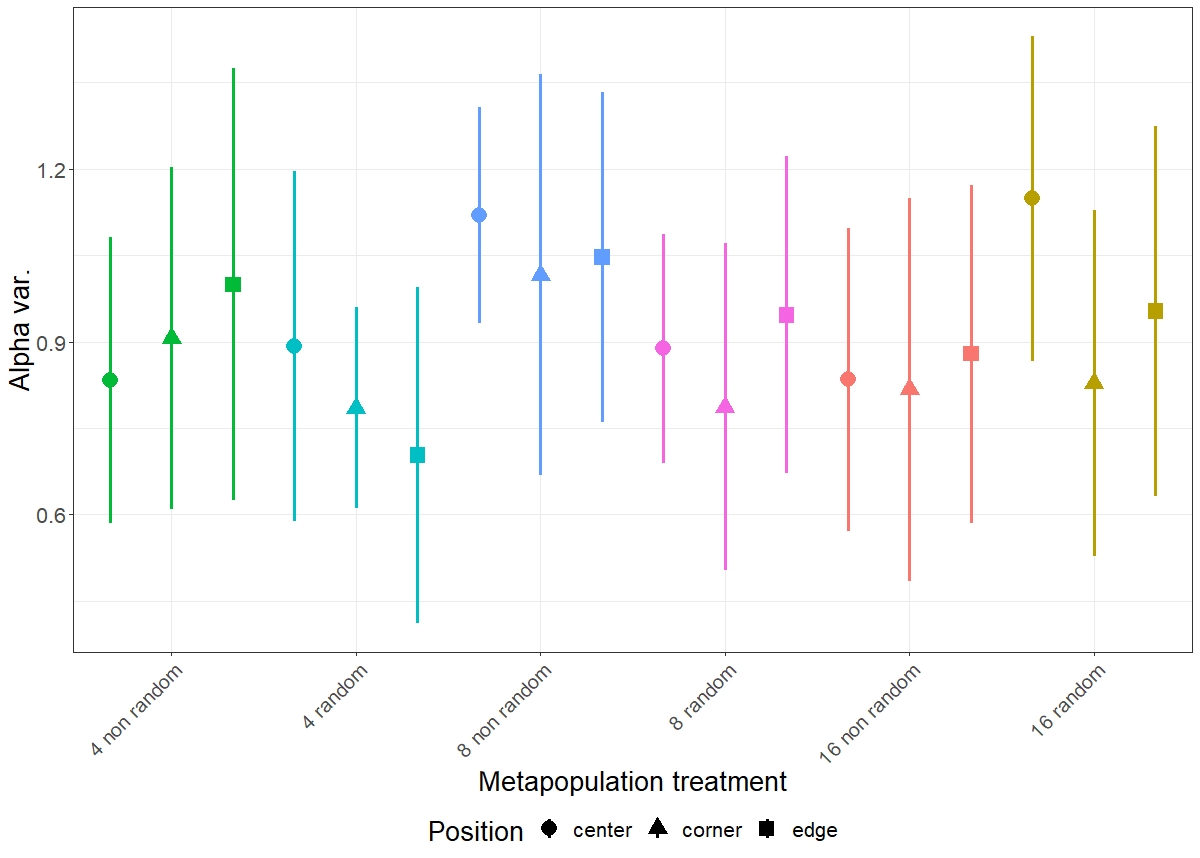

***Table SI2.2a***

| **Explanatory variable** | **Χ^2^** | **Df** | **Pr(>Χ^2^)** |
| --- | --- | --- | --- |
| Global connectedness | 3.7361 | 2 | 0.1544 |
| Randomization | 0.0017 | 1 | 0.9674 |
| Local connectedness | 2.0694 | 2 | 0.3553 |
| Global connectedness * Randomization | 3.7475 | 2 | 0.1535 |
| Global connectedness*Local connectedness | 0.3226 | 4 | 0.9883 |
| Randomization*Local connectedness | 1.5933 | 2 | 0.4508 |
| Global connectedness*Randomization*Local connectedness | 1.9433 | 4 | 0.7462 |

***Table SI2.2b***

| **Explanatory variable** | **Χ^2^** | **Df** | **Pr(>Χ^2^)** |
| --- | --- | --- | --- |
| Global connectedness | 2.9005 | 2 | 0.2345 |
| Randomization | 0.0693 | 1 | 0.7923 |
| Local connectedness | 1.8493 | 2 | 0.3967 |
| Global connectedness * Randomization | 2.9729 | 2 | 0.2262 |
| Global connectedness*Local connectedness | 1.4194 | 4 | 0.8408 |
| Randomization*Local connectedness | 2.8874 | 2 | 0.2360 |
| Global connectedness*Randomization*Local connectedness | 3.9040 | 4 0 | 0.4191 |

*Table SI2.2: ANOVA (Analysis of Variance: Wald Type-III chisquare test) results for population size (a) and alpha variability (b) of the local populations over 4 weeks before the end of the experimental evolution.*

**Supplementary Information 3: dispersal tendency and starting density**

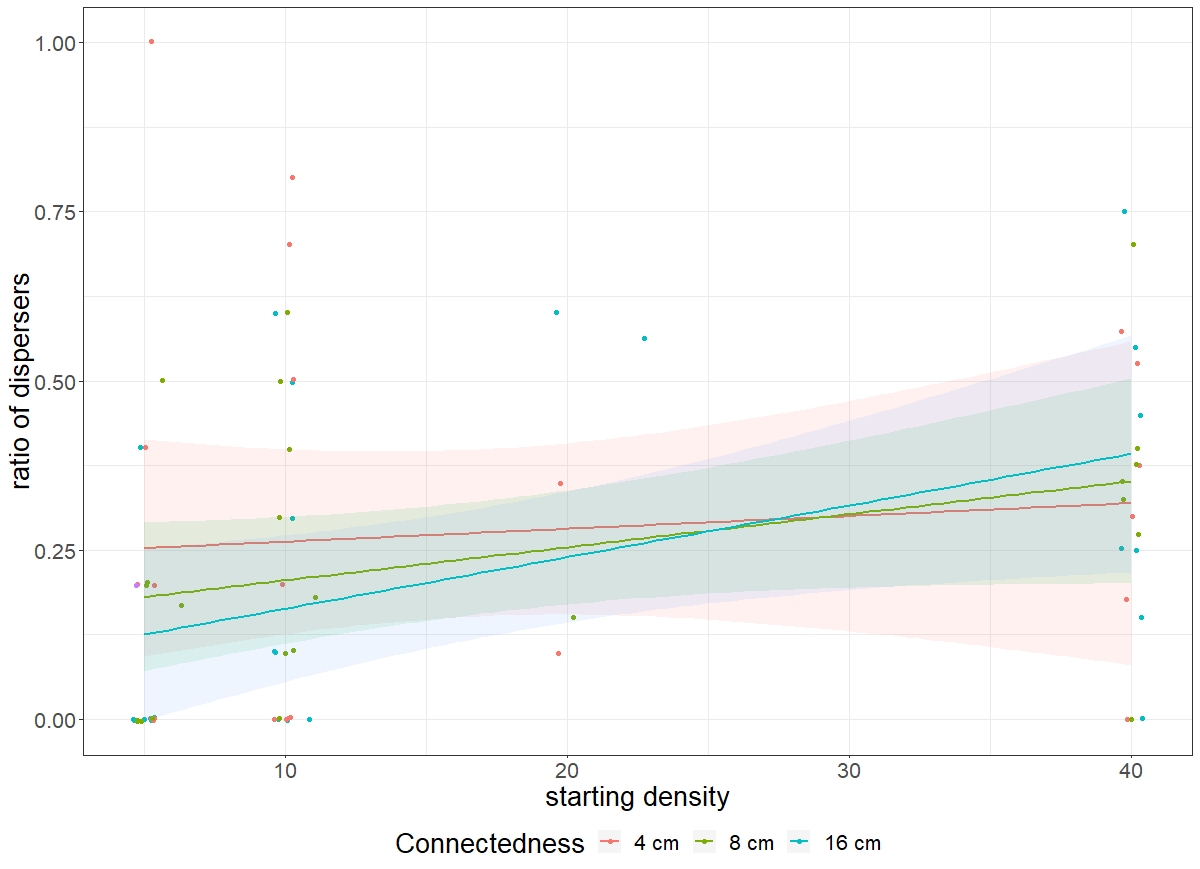
*Figure SI3.1: effect of starting density (individuals per patch) over the total dispersal rate after 4 days. Points were jittered on the x-axis to avoid superimpositions*.

**Supplementary Information 4: reproductive performance test**

1) *Days of death*

| **Explanatory variable** | **χ^2^** | **df** | **Pr(>χ^2^)** |
| --- | --- | --- | --- |
| Position | 4.8999 | 3 | 0.1793 |
| Fragmentation | 3.1986 | 2 | 0.2020 |
| Randomization | 2.1091 | 1 | 0.1464 |
| Position*Fragmentation | 3.1921 | 4 | 0.5262 |
| Position*Randomization | 1.8945 | 2 | 0.3878 |
| Fragmentation*Randomization | 3.6656 | 2 | 0.1600 |
| Position* Fragmentation*Randomization | 3.2045 | 4 | 0.5242 |

*Table SI4.1: ANOVA (Analysis of Variance: Wald Type-III chisquare test) results for the day of death of females during the reproductive performance test.*

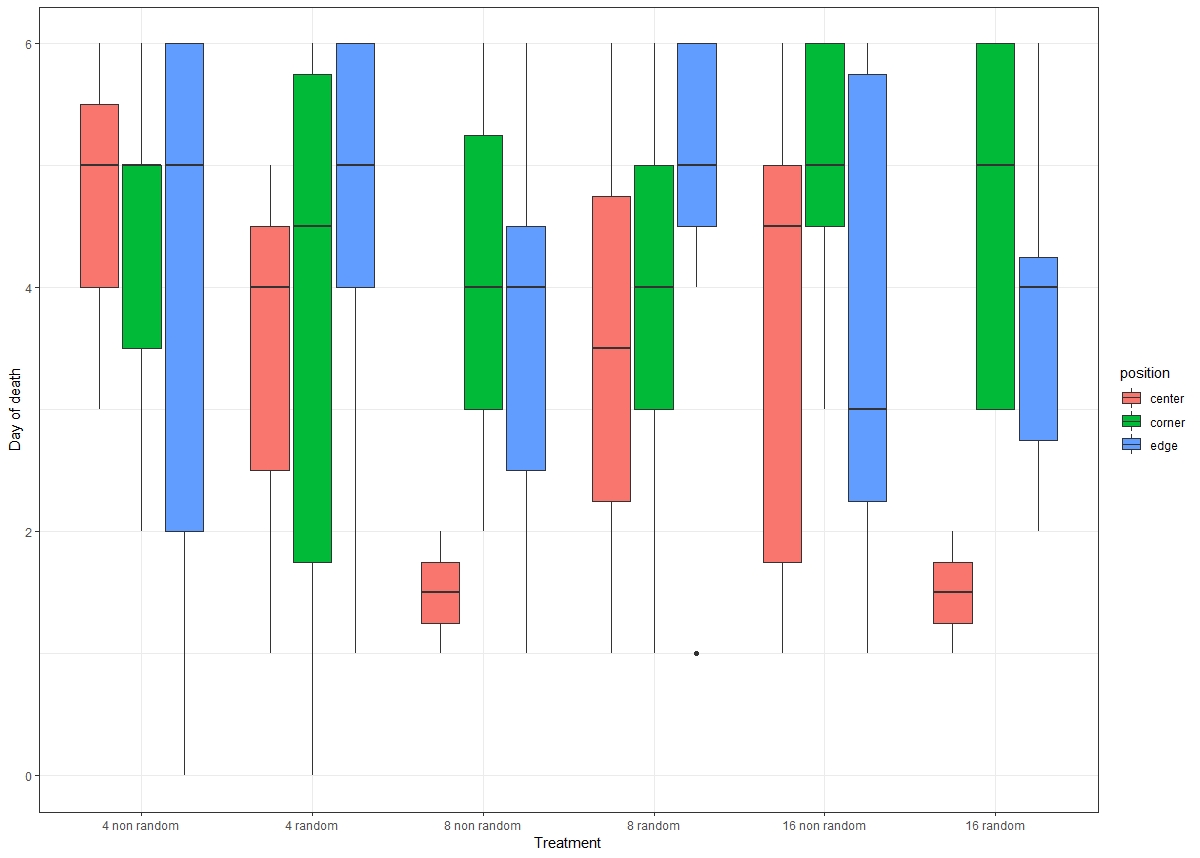
*Fig. SI4.1: Day of death for F1 females during the reproductive performances test*

2) *Reproductive performances*

| **Explanatory variable** | **χ^2^** | **df** | **Pr(>χ^2^)** |
| --- | --- | --- | --- |
| Local Connectedness | 9.6736 | 2 | 0.007933(**) |
| Metapopulation Connectedness | 3.2587 | 2 | 0.196056 |
| Randomization | 0.4209 | 1 | 0.516487 |
| Local Connectedness * Metapopulation Connectedness | 1.6079 | 4 | 0.806379 |
| Local Connectedness * Randomization | 0.8291 | 2 | 0.660647 |
| Metapopulation Connectedness * Randomization | 3.7884 | 2 | 0.150437 |
| Local Connectedness * Metapopulation Connectedness * Randomization | 1.1641 | 4 | 0.883968 |

*Table SI4.2: Full Model Analysis of Deviance table (Wald χ² test) for intrinsic reproductive performances as determined by the number of offspring produced by a single female after 6 days.*

**Supplementary Information 5: Correlation analysis**

a) ***Pearson’s correlation coefficients***

|  | **Reproductive Performance** | **Starvation Resistance** | **Mean population size** | **Dispersal timing** | **Alpha Variability** |
| --- | --- | --- | --- | --- | --- |
| **Reproductive Performance** | - | -0.11920059 | 0.88301079 | 0.5176116 | -0.22682566 |
| **Starvation Resistance** | *0.5880* | - | 0.04188302 | -0.1618848 | -0.06601743 |
| **Mean population size** | *0.0000* | *0.8495* | - | 0.4259799 | -0.16379561 |
| **Dispersal**  **timing** | *0.0114* | *0.4605* | *0.0427* | - | -0.29319649 |
| **Alpha**  **Variability** | *0.2980* | *0.7647* | *0.4552* | *0.1745* | - |

*Table SI5.1: Pearson’s correlation coefficients (above diagonal) and significance values (below diagonal, italic) between evolved traits, local population size and variability over the 6 months of the experimental evolution.*
